# You can do it!–Using published undergraduate research on *Hydra* mouth opening to train undergraduates

**DOI:** 10.1101/2024.12.05.627060

**Authors:** S. Hackler, T. Goel, J. Pacis, E.-M. S. Collins

## Abstract

Biophysical research is both exciting and challenging. It is exciting because physical approaches to biology can provide novel insights, and it is challenging because it requires knowledge and skills from multiple disciplines. We have developed an undergraduate biophysical laboratory module that is accessible to both biology and physics majors, teaches fundamental skills such as time-lapse microscopy, image analysis, programming, critical reading of scientific literature, and basics of scientific writing and peer-review. This module uses published research on the biomechanics of *Hydra* mouth opening as its framework because this work was co-first authored by an undergraduate student and featured in the public press, thus providing two anchors that make this research accessible and exciting to undergraduates. Students start with a critical reading and discussion of the publication. Then they execute some of the experiments and analysis from the paper, thereby learning fluorescence time-lapse microscopy and image analysis using ImageJ and/or MATLAB/Python, and compare their data to the literature. The module culminates in the students writing a short paper about their results following the *Micropublication* journal style, a blinded peer-review, and final paper submission. Here, we describe one possible implementation of this module with the necessary resources to reproduce it, and summarize student feedback from a pilot run. We also provide suggestions for more advanced exercises. Several students expressed that repeating a published study done by an undergraduate student inspired and motivated them, thus creating buy-in and assurance that they “can do it”, which we expect to help with confidence and retention.

## I. Introduction

The 2022 decadal report on the Physics of Living Systems published by the National Academies of Sciences, Engineering, and Medicine identified that, despite two decades of growth in the number of doctoral degrees awarded for biophysics, the subject remains under-represented in undergraduate curricula (National Academies of Sciences, 2022). The report calls on both physics and biology departments to integrate the Physics of Living Systems into their courses (National Academies of Sciences, 2022). While undergraduate biology curricula have seen major changes (Guenther et al., 2019; Wei & Woodin, 2011) in response to the recommendations of the 2009 “Vision and Change: A Call to Action” report (American Association for the Advancement of Science, 2009), to focus on the mastery of core biological principles and skills, biophysics is not an integral part of the curriculum. Similarly, while introductory physics courses for non-physics majors have been reformed to include more life science applications, which has been found to increase engagement of biology majors with physics (see e.g. (Geller et al., 2018)), upper-level biophysics courses that build upon the introductory course material are sparse. Thus, there is a need and opportunity for biophysics educators to develop upper-level undergraduate biophysics courses that appeal to both physics and biology departments, incorporate pedagogical reforms, and are accessible to students from diverse backgrounds.

In both physics and biology, pedagogical reforms like inquiry-based labs have been shown to broadly benefit student learning, improve students’ attitudes towards learning science (Brownell et al., 2015; Geller et al., 2018; Jeffery et al., 2016; Wilcox & Lewandowski, 2017) and increase students’ performance in data analysis and interpretation (Brownell et al., 2015; Karelina et al., 2007). Course-Based Undergraduate Research Experiences (CUREs) (Auchincloss et al., 2014) have been shown to increase confidence and student persistence through a STEM degree (Graham et al., 2013), especially for students coming from underrepresented backgrounds (Bangera & Brownell, 2014; Bradshaw et al., 2023). Research has also shown that students who begin the semester with weaker experimental design skills show greater gains than initially higher-performing students (Blumer & Beck, 2019). Other important aspects of incorporating research into undergraduate education have emphasized advanced laboratory techniques (Full et al., 2015), reading and discussing research papers (Parent et al., 2010), and laboratory modules based on ongoing research by faculty at the instructional institution (Van Hecke et al., 2002). Moreover, it has been emphasized that student buy-in, metacognition, and an understanding of learning goals are critical for improving students’ *perception* of their career readiness and persistence in STEM (Etkina & Murthy, 2005).

Here, we introduce a multi-week, research-based laboratory module that was developed for an intermediate level systems biology course and incorporates pedagogical elements that address student buy-in and skill development. Its content is based on a published biophysics article in the field of tissue mechanics (Carter et al., 2016) and is accessible to both physics and biology undergraduate students. The first author of the published work was an undergraduate student, and this interdisciplinary lab was designed to provide students a parallel experience to that described in the article. Because the paper in question was authored by an undergraduate student, students feel empowered that they can do similar work and approach the lab module with more confidence. This increases student buy-in and resilience considering doing technically challenging experimental and computational tasks. The module provides a unique opportunity for undergraduates to experience the research process by performing published experiments, analyzing their own data rigorously and authentically, reconciling their findings with existing literature, and formally reviewing and presenting their work.

We chose to examine mouth opening in *Hydra* for several reasons:

- In understanding the underlying mechanism for *Hydra* mouth opening, students are exposed to cellular and organismal biology, as well as to the biomechanics of soft tissues. Thus, the background preparation requires the discussion of both biological and physical principles, which immediately invites students to draw from multiple disciplines when working on this module.
- The interdisciplinarity of the system allows for multiple avenues of analysis, enabling the instructor to customize the task to meet the needs and interests of their students. Due to the wide variability in students’ comfort with computer programming, which can dissuade some students from engaging with the computational analysis featured in the module, this flexibility is essential for managing student confidence and maximizing buy-in.
- The experimental techniques are within reach of most students with previous biology laboratory experience.
- *Hydra* are easy to culture in the lab and can be multiplied by bisecting them and allowing the pieces to regenerate. With the availability of fluorescently labelled transgenic lines (Juliano et al., 2014; Wang et al., 2022; Wittlieb et al., 2006) and reversible anesthetics (Goel et al., 2019), *Hydra* are easy to image and to manipulate surgically. They are also easy to manipulate pharmacologically due to their small size and aquatic environment. For example, in the absence of prey, mouth opening can be induced using reduced glutathione (Carter et al., 2016) and quinine hydrochloride (Goel et al., 2024).
- The software needed to analyze the rate of mouth opening is freely available, requires minimal technical expertise, and is widely used in professional research settings across multiple disciplines.
- There are many optional avenues for increasing the complexity of the module to accommodate students with more or less experience with computational techniques and image analysis, which we will discuss later.
- The lab module lends itself well to a remote classroom if needed – in this case, the instructor can share the sample data provided in the Supplemental Material with their students.
- Because the original work was authored by an undergraduate researcher, students can get inspired as they can identify. It gives them confidence to know that an undergraduate student like them did it before.

The four key components of the lab module address basic training in the sciences that all undergraduate students majoring in the natural sciences should obtain: 1) How to critically read a scientific paper, 2) How to plan and perform an experiment and reconcile results with the published literature, 3) How to interpret and present results, and 4) How to write and peer review scientific papers.

## II. Scientific and Pedagogical Background

### A. Scientific Background – The Biomechanics of *Hydra* mouth opening

*Hydra* are cnidarian polyps found in freshwater sources around the globe. *Hydra* have a cylindrical body column that is a few hundred microns in diameter and a couple of centimeters in length (Vogg et al., 2019). At one end of the body column is an adhesive foot which *Hydra* uses to stick to substrates. At the other end is the head, consisting of a conical structure called the hypostome surrounded by a ring of tentacles (**Figure 1A**). The tentacles allow *Hydra* to catch and incapacitate prey using specialized cells called nematocytes. The prey is then moved toward the mouth at the apex of the hypostome and ingested. *Hydra* consists of two epithelial layers, an outer ectoderm and an inner endoderm, connected to each other by an extracellular matrix called the mesoglea (Vogg et al., 2019). Both layers also contain specialized cells, which include neurons (both layers), gland cells (endoderm), and nematocytes (primarily ectoderm) (Bode, 2009). The epitheliomuscular cells of the ecto- and the endoderm contain 1-2 cell diameter short extracellular actin projections called myonemes that can generate contractile forces (Davis, 1974) (**Figure 1C**). These myonemes appear arranged as radial spokes that originate in the center of the hypostome in the ectoderm and as concentric circles in the endoderm when looking top-down onto the head (**Figure 1B**). Embedded in each epithelial layer is a neuronal network which receives and transmits environmental signals and coordinates behaviors (Dupre & Yuste, 2017).

**Figure 1:**
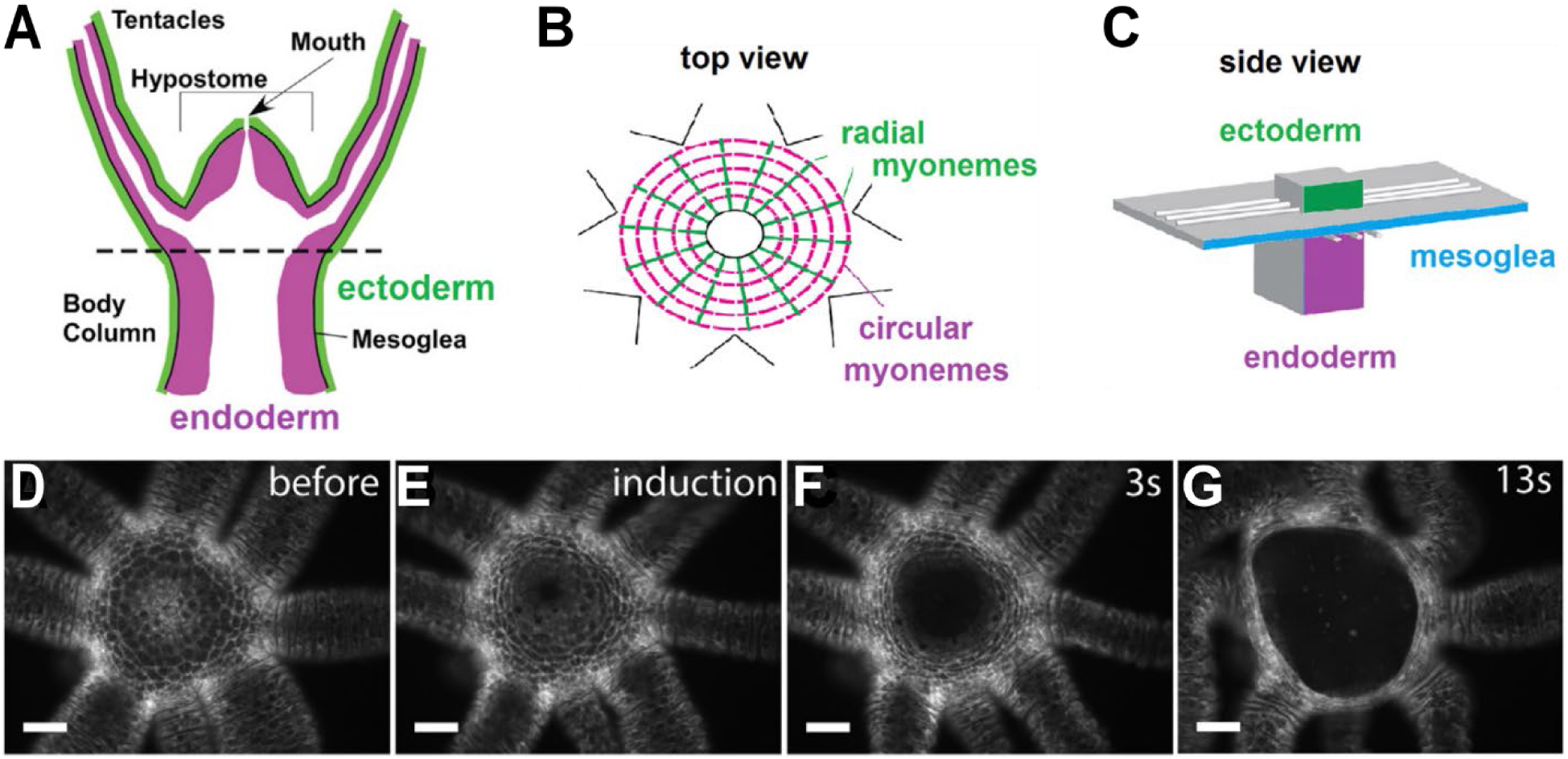
Overview of *Hydra* anatomy and mouth opening. (A) Sideview schematic of the *Hydra* head showing the hypostome and tentacles and the two epithelial layers. (B) Top-view schematic of the *Hydra* head showing myoneme arrangement in the ectoderm (green) and endoderm (magenta) epithelial layers. (C) Side-view schematic showing the perpendicular orientation of myonemes in the endoderm and ectoderm on a cell-level. (D-G) Example fluorescence microscopy images of a chemically-induced mouth opening event. The ectodermal cell layer is visible. Scale: 100 microns. Figure modified from Carter et al. (Carter et al., 2016) with permission from authors.

Following neuronal activation due to environmental chemical signals (such as food or certain chemicals, including reduced glutathione and quinine hydrochloride (Forrest, 1962; Kulkarni & Galande, 2014; Lenhoff, 1961)), the ectodermal myonemes in the hypostome contract, causing a hole (“the mouth”) to form at the apex of the hypostome (**Figure 1D-G**). This mouth opening widens over the course of a few seconds, sometimes exceeding the diameter of the body column (Carter et al., 2016). After prey has been ingested, the mouth re-seals. When waste gets expelled or osmotic pressure needs to be released, the mouth re-opens. The forces necessary to create the mouth opening are generated by myonemes that act at short range over the millisecond timescale but produce a large symmetric tissue deformation over the seconds timescale, in the absence of centralized neuronal or chemical coordination (Goel et al., 2024). Mouth opening does not involve any cellular rearrangements – the tissue deformation is accomplished by shape changes of the epithelial cells (Carter et al., 2016). Thus, mouth opening is a fascinating problem to study – from the biological perspective of control and coordination of behavior, and from the physics perspective of large deformations of soft tissue arising from short range, uncoordinated forces.

### B. Pedagogical Background

This 6-week laboratory module was developed for a semester long course in systems biology (max enrollment, n=24) taught at Swarthmore College, which is a highly selective small residential liberal arts college in the suburbs of Philadelphia, in Pennsylvania. In 2022/23, when this lab module was taught, the self-reported ethnic and racial identity of the student body at the college was 32% White, 18% Asian, 14% Hispanic, 10% Bi-Racial (non-Hispanic), and 9% African-American (Swarthmore College, 2024) Students enrolling in systems biology need to meet a few prerequisites, including having taken an introductory biology course in molecular and cellular biology (or have AP credit), an introductory statistics course, and differential calculus.

Historically, students enrolling in the course come from different disciplines and with different levels of preparation in relevant STEM fields. During the semester in which this module was first implemented, 15 students were enrolled in the course, with the majority being second year biology majors (**Figure 2A, B**). Coming into the course, students reported a high level of proficiency in biology, moderate math proficiency, and relatively low proficiencies in physics and programming (**Figure 2C**).

**Figure 2:**
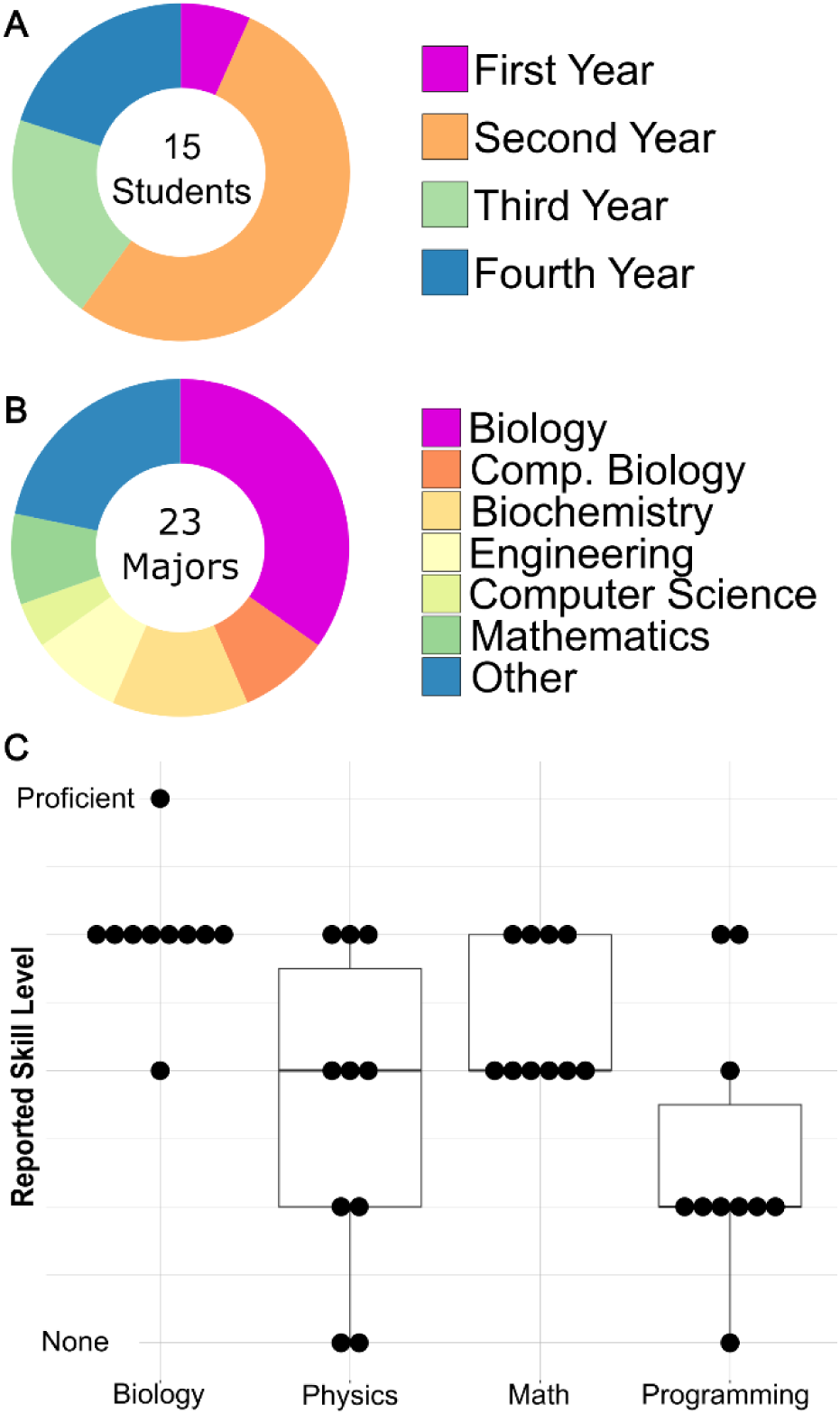
Student demographics and preparation. Breakdown of enrolled students by (A) class year and (B) declared major. Double majors are individually counted, so the total number of majors represented (23) exceeds the enrollment in the course (15). Of the 15 students, only two were not majoring in Biology, Computational Biology, or Biochemistry. The “Other” category includes Spanish, English literature, Art, Economics, and Architecture majors. (C) Student’s self-reported proficiency with relevant subject areas before the *Hydra* mouth opening module (n=12). The vertical axis represents each student’s self-assessed proficiency in the given subject area. Each point represents a single student and the boxes span the 25^th^ and 75^th^ percentile responses.

The lab module was offered in two sections: section A was aimed at students with prior computational experience and section B was for students without said experience. Students could self-select during enrollment but the faculty member teaching the course would double-check their pre-requisites to ensure appropriate placement. Each lab section was provided with the same course materials, allowing students who enrolled in section A to follow the program of section B if they discovered that they were not comfortable with programming on their own. Each lab section was taught by the faculty member and a professional laboratory instructor. Each section met once weekly for 3 h and 15 min. Enrollment in either section was limited to a maximum of 12 students. Students worked in pairs at designated lab stations, which were equipped as necessary for the lab module. Because the total enrollment in spring 2023 was 15 students, one group consisted of 3 students.

Since most of the students had taken at least one introductory biology course at the college, they had a basic understanding of laboratory safety. However, because some individuals had not completed biology laboratory safety training, a brief module-specific training was provided, to ensure safe working conditions. This was important as this module involves working with chemicals, sharps, and biohazardous waste.

An anonymous survey (included as Supplemental Material and reviewed and approved by the Swarthmore College Institutional Review Board (IRB-FY24-25-19)) was administered post-course completion to collect students’ feedback on the module. The response rate was 67%.

### C. Module Framework

The overarching goal of this lab module is to provide students with an authentic experience of interdisciplinary research. To that end, this module aims to improve student aptitude for performing research with live biological samples, using computation to analyze images from fluorescent microscopy, and reading, writing, and discussing scientific papers. Instead of having an inquiry-based lab module wherein students conduct original research, we chose to base our module on an undergraduate student first authored paper. The publication we chose on the biomechanics of *Hydra* mouth opening by Carter et al. (Carter et al., 2016) has multiple advantages that make it well-suited for inspiring undergraduate students: 1) The paper’s first author is an undergraduate student, thus showing students that they could do research like this, 2) The experiments are visually pleasing and have been widely featured in the public press, such as in the “How stuff works” video series on YouTube (HowStuffWorks, 2016), thus bringing in an excitement factor to get students interested, and 3) It is fairly easy to read without requiring a lot of background knowledge in the field, allowing students to connect without having to do much additional reading. To prepare students for the group discussion of the Carter et al paper, the instructor should provide them with some introductory material on mechanics and biology, depending on their level of background preparation. Though reproducing published research sacrifices some of the freedom of inquiry-based lab modules, it grounds the experience in the context of published research and encourages sophisticated reflection and discussion throughout the module, by being able to compare one’s data with the published work. There are three key components that we believe are necessary for the success of this kind of laboratory module:

#### Framing and Metacognition

Students must be encouraged to think about how they might have conducted the research in the paper from the start. This includes engaging students in thinking about not only the context and the science behind the experiments but also the logistics of the experiment, their ability to analyze the data and ways of communicating their findings through a scientific paper. Students are asked to read the Carter et al. paper in preparation for this module and answer a set of reading reflection questions.

The content reflection questions (**Table 1**) asked students to engage with the scientific ideas presented in the paper. The personal reflection questions asked students to consider their own abilities and confidence in writing a research paper and to brainstorm how they might overcome some of the elements that they might find especially challenging. These reflections are initially individual but are then incorporated into small group discussions and eventually a class-wide discussion. Beginning the entire module with these kinds of prompts encourages students to consider the challenges of writing and publishing academic research and identify the specific skills that they need to develop to prepare themselves for a research career.

**Table 1:** Reading reflection questions.

| **Content reflection** | **Personal reflection** |
| --- | --- |
| What were the most significant results/findings? | Knowing your strengths and weaknesses, which parts would be easy for you to write? Which would be especially difficult? Why? |
| Do you agree with the interpretation of the key results? Why or why not? | Can you think of strategies to help you write the difficult parts more effectively? |
| How did this paper advance the field? | Can you think of strategies to help you write the difficult parts more effectively? |

#### Experimental Techniques

The techniques required for data collection and analysis are examples of current and transferrable skills that may serve students in their future academic and professional lives. This module prepares students to work with aquatic invertebrates, use fluorescence microscopy and video capture, and take measurements from recorded images and video using computational image analysis. Students need to be given ample time to independently engage with every step of the research process to ensure continuity from literature review to data collection and analysis, and eventually, to publication. In the case of *Hydra* mouth opening, this means providing students with the time, materials, and instructions beginning with intact animals, guiding them through sample preparation and image capture, and outlining the necessary steps of image processing (all described in the following section). In addition to ensuring continuity in the students’ experience, this approach teaches students skills and techniques that are commonplace in professional research labs and provides numerous jumping off points where motivated or advanced students could extend their analysis to examine the system in more sophisticated ways.

#### Presentation

Students are asked to present their findings and interpretations in the form of a lab report mirroring a scientific publication. We used the short paper format of *MicroPublication* (https://www.micropublication.org/) because of the limited time available for the module (6 weeks) and because this course is not a formal writing course. Students were provided with example papers from this publisher to have a framework for their own writing and general guidelines about scientific writing and figure making. One could easily expand the writing portion of this lab and follow a different publication format and extend it into a more intensive, multi-week experience. An important feature of the writing is that students give and receive peer reviews on their reports, to mimic the process of scientific publication. Students then get to incorporate reviewer feedback in generating their final reports. To incentivize students to make thoughtful and serious comments on the reports they review, a small portion of each student’s grade for the module is based on the level of detail and thoughtfulness of the comments they produce for their peers. This process, of discussion, drafting, feedback, and revision, is at the core of the endeavor to publish scientific research.

## III. Materials and Methods

After students have had a chance to engage with the primary literature and discussed the experimental protocol, it is useful to break the experiment down into stages, including sample preparation, mounting samples for imaging, and image acquisition and analysis. The protocols for each of these stages and pointers toward additional resources are provided in the lab handout (Supplemental Material) and the required materials for implementation are listed in **Table 2**.

**Table 2:**
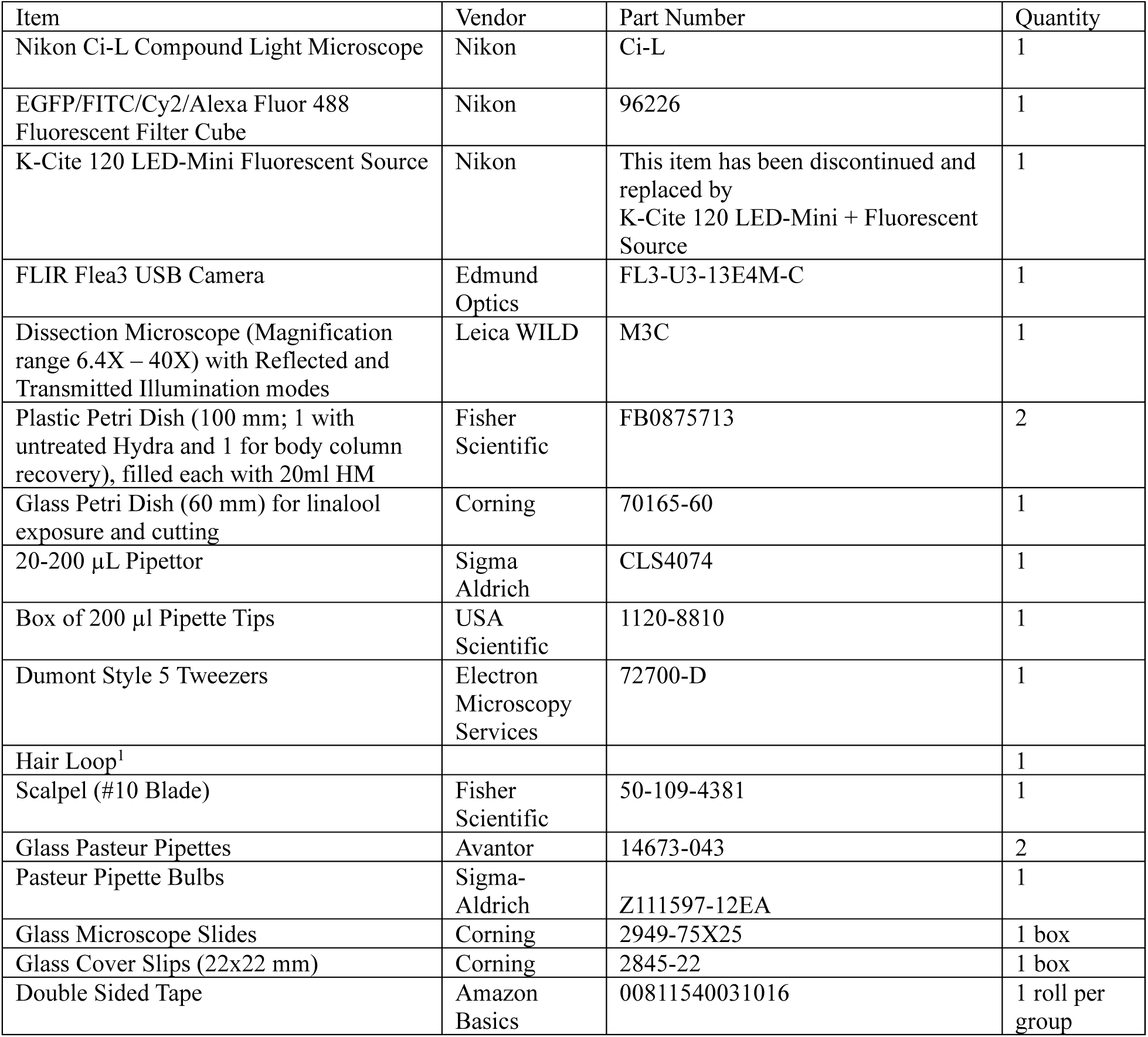
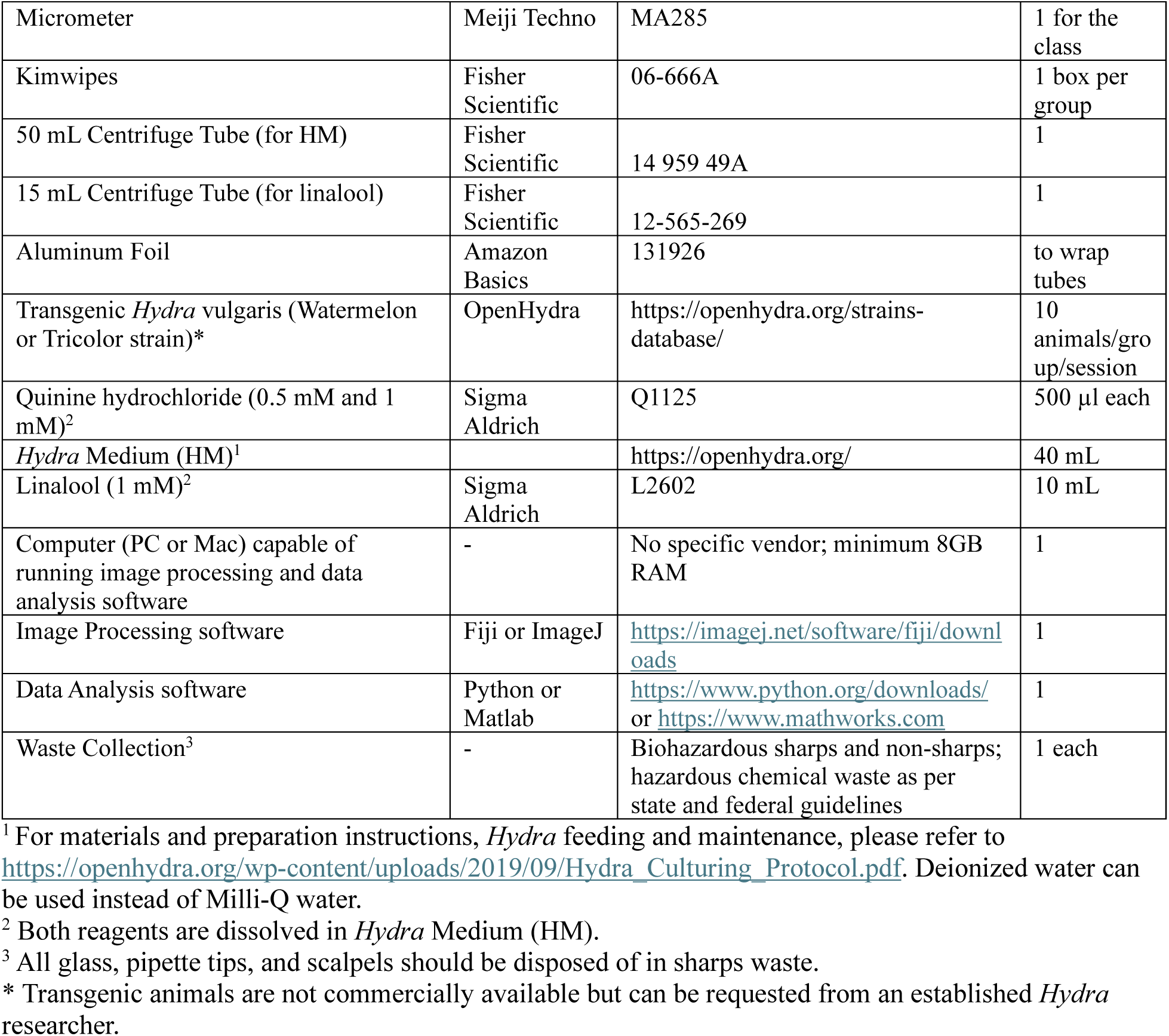
Overview of components used in the pilot implementation of the *Hydra* lab module. Equivalent items from other vendors can be used instead. The indicated quantities are listed for a single experimental station. In our case, students worked in pairs at each station.

| Item | Vendor | Part Number | Quantity |
| --- | --- | --- | --- |
| Nikon Ci-L Compound Light Microscope | Nikon | Ci-L | 1 |
| EGFP/FITC/Cy2/Alexa Fluor 488 Fluorescent Filter Cube | Nikon | 96226 | 1 |
| K-Cite 120 LED-Mini Fluorescent Source | Nikon | This item has been discontinued and replaced by K-Cite 120 LED-Mini + Fluorescent Source | 1 |
| FLIR Flea3 USB Camera | Edmund Optics | FL3-U3-13E4M-C | 1 |
| Dissection Microscope (Magnification range 6.4X – 40X) with Reflected and Transmitted Illumination modes | Leica WILD | M3C | 1 |
| Plastic Petri Dish (100 mm; 1 with untreated Hydra and 1 for body column recovery), filled each with 20ml HM | Fisher Scientific | FB0875713 | 2 |
| Glass Petri Dish (60 mm) for linalool exposure and cutting | Corning | 70165-60 | 1 |
| 20-200 μL Pipettor | Sigma Aldrich | CLS4074 | 1 |
| Box of 200 μl Pipette Tips | USA Scientific | 1120-8810 | 1 |
| Dumont Style 5 Tweezers | Electron Microscopy Services | 72700-D | 1 |
| Hair Loop<sup>1</sup> |  |  | 1 |
| Scalpel (#10 Blade) | Fisher Scientific | 50-109-4381 | 1 |
| Glass Pasteur Pipettes | Avantor | 14673-043 | 2 |
| Pasteur Pipette Bulbs | Sigma-Aldrich | Z111597-12EA | 1 |
| Glass Microscope Slides | Corning | 2949-75X25 | 1 box |
| Glass Cover Slips (22x22 mm) | Corning | 2845-22 | 1 box |
| Double Sided Tape | Amazon Basics | 00811540031016 | 1 roll per group |
| Micrometer | Meiji Techno | MA285 | 1 for the class |
| Kimwipes | Fisher Scientific | 06-666A | 1 box per group |
| 50 mL Centrifuge Tube (for HM) | Fisher Scientific | 14 959 49A | 1 |
| 15 mL Centrifuge Tube (for linalool) | Fisher Scientific | 12-565-269 | 1 |
| Aluminum Foil | Amazon Basics | 131926 | to wrap tubes |
| Transgenic *Hydra* vulgaris (Watermelon or Tricolor strain)* | OpenHydra | https://openhydra.org/strains-database/ | 10 animals/group/session |
| Quinine hydrochloride (0.5 mM and 1 mM)<sup>2</sup> | Sigma Aldrich | Q1125 | 500 µl each |
| *Hydra* Medium (HM)<sup>1</sup> |  | https://openhydra.org/ | 40 mL |
| Linalool (1 mM)<sup>2</sup> | Sigma Aldrich | L2602 | 10 mL |
| Computer (PC or Mac) capable of running image processing and data analysis software | - | No specific vendor; minimum 8GB RAM | 1 |
| Image Processing software | Fiji or ImageJ | https://imagej.net/software/fiji/downloads | 1 |
| Data Analysis software | Python or Matlab | https://www.python.org/downloads/ or https://www.mathworks.com | 1 |
| Waste Collection<sup>3</sup> | - | Biohazardous sharps and non-sharps; hazardous chemical waste as per state and federal guidelines | 1 each |

<sup>1</sup> For materials and preparation instructions, *Hydra* feeding and maintenance, please refer to https://openhydra.org/wp-content/uploads/2019/09/Hydra_Culturing_Protocol.pdf. Deionized water can be used instead of Milli-Q water.
<sup>2</sup> Both reagents are dissolved in *Hydra* Medium (HM).
<sup>3</sup> All glass, pipette tips, and scalpels should be disposed of in sharps waste.
\* Transgenic animals are not commercially available but can be requested from an established *Hydra* researcher.

Below, we emphasize steps from the experiment that present learning opportunities for students and highlight possible modifications to suit specific teaching contexts.

### Sample preparation

Students were provided transgenic ‘watermelon’ (WM) (Glauber et al., 2013) or tricolor (Wang et al., 2022) *Hydra* and the materials necessary for sample preparation (**Table 2**) at their workstation. 1mM linalool (anesthetic solution) (Goel et al., 2019) and 0.5mM and 1mM quinine hydrochloride (stimulants that causes mouth opening) (Goel et al., 2024) solutions were also provided to students. Students were required to read the safety data sheet (SDS) and protocols for usage for these chemicals prior to starting any experiments. Both chemicals are light sensitive and must be kept away from light. Both chemicals are combustible and skin irritants. It is therefore important to wear proper personal protective equipment while executing the experiments and disposing of unused chemicals and contaminated solid materials in the appropriate hazardous waste containers. This is an opportunity to explain to students why it is indispensable to read protocols and material safety data sheets as part of the lab preparation. Transgenic *Hydra* are considered biohazardous materials and need to be handled and discarded following state and federal regulations.

Students used linalool to anesthetize the *Hydra* before decapitating them and then waited for them to recover from anesthesia before they mounted their heads for mouth opening experiments. The decapitation step can also be done without linalool treatment. However, the linalool treatment relaxes the *Hydra,* making it easier to obtain head samples without excessive body column tissue. Too much body column tissue makes it difficult to orient the *Hydra* head for top-down imaging. It is difficult to know when linalool has taken effect and when it has worn off by simply looking at the animals. Because body columns react differently to being pinched by a pair of forceps when they are anesthetized compared to when they are not (**Figure 3A**), the “pinch response” test can be used to evaluate the anesthesia. Untreated and fully recovered body columns contract globally in response to a pinch at the lower part of the body column whereas anesthetized *Hydra* contract only locally (**Figure 3A**) (Goel et al., 2019). Students should be encouraged to recognize that if they did not have the pinch test on the intact *Hydra* as a readout, it would be hard to determine when the effect of the linalool had worn off and that linalool may affect mouth opening. Thus, this is an opportunity to teach students about the need of designing appropriate controls/markers for whatever treatment they use in experiments.

**Figure 3.**
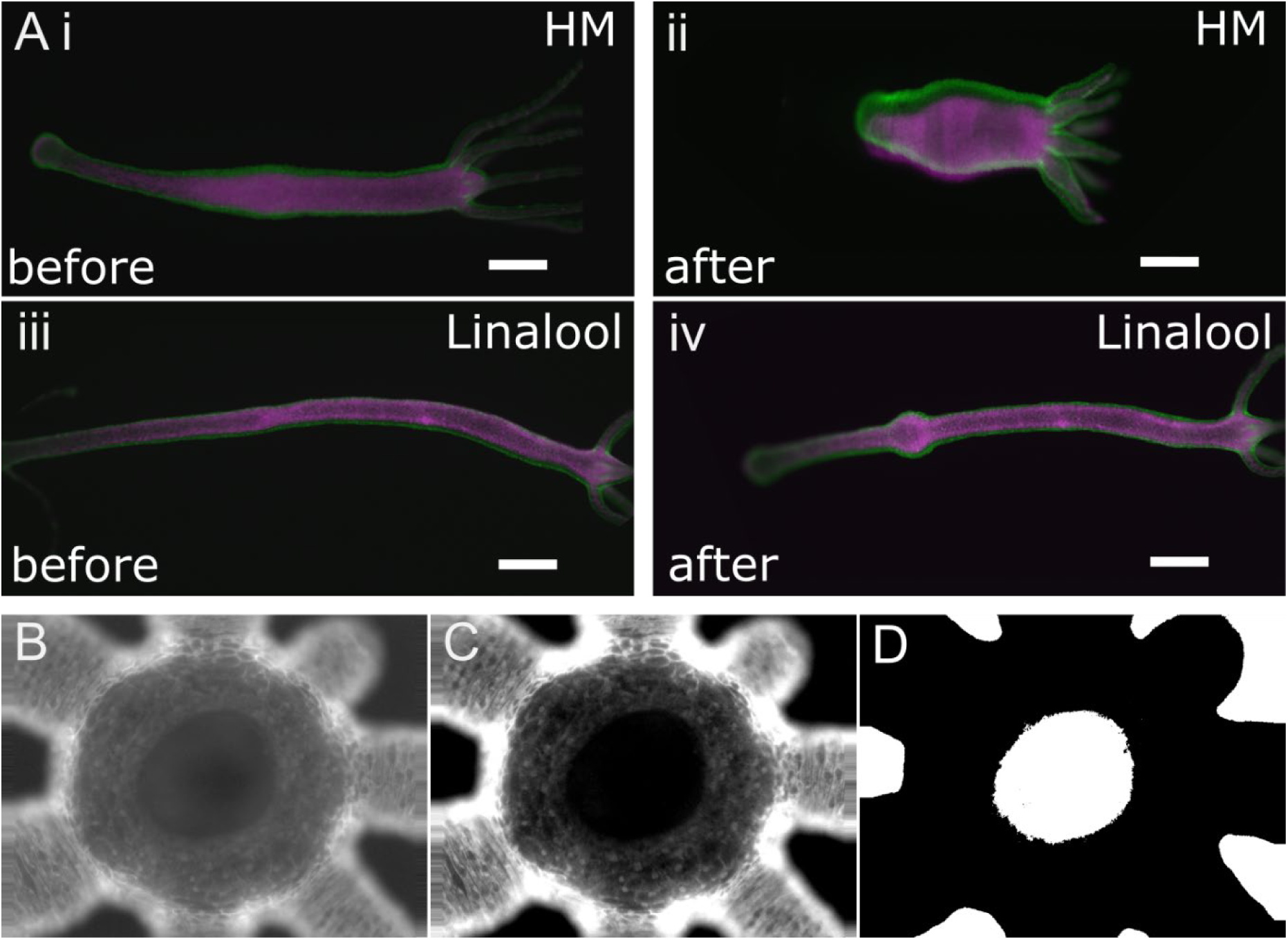
Methods for studying mouth opening. (A) The *Hydra* pinch response is used to test when the anesthetic (linalool) is effective or has worn off. (i) An untreated, elongated *Hydra* before pinching with forceps near the foot. (ii) The animal contracts when pinched with forceps. Due to the time delay from filter switching (0.6 seconds), there is a slight mismatch of the tissue layers. (iii-iv) A *Hydra* incubated in 1 mM linalool for 15min before (iii) and after (iv) pinching near the foot. The pinch-induced contraction is local (bulging) and no global contraction is observed. Scale bar: 300 microns. (C-E) Illustration of the basic image analysis steps to isolate the mouth from the image using image contrast enhancement and thresholding. (B) Raw data, (C) contrast enhanced image, (D) thresholded image.

### Data collection

Students were given a demonstration on how to use the fluorescence microscope and collect data using a camera mounted on the microscope by one of the instructors.

Depending on the level of familiarity students have with fluorescence microscopy, instructors can decide what level of details to share with the students about the physics behind fluorescence, how it used as a tool in biology and the optics behind fluorescence imaging. As students collect data, this is a good opportunity to discuss the tradeoff between collecting data at high temporal resolution versus the storage space available. There are two limitations here – the physical storage available to students and the computing power available to process large single videos at a time. It is also important to emphasize how imaging the micrometer (or some object of known size) is important to get the scale factor of the microscope at the given magnification.

Students in both lab sections were also given a tutorial on how to use Image J to extract the *Hydra* mouth areas from the image data (**Figure 3B-D**), plot the mouth area as a function of time in MATLAB, and how to use the MATLAB curve fitting toolbox to fit the mouth area curves to the logistic model (equation 1) provided in the Carter paper (Carter et al., 2016). Students with prior computational experience in section A were provided with the main steps for analyzing the images but were expected to write their own code to do so. However, all students were introduced to ImageJ (Schindelin et al., 2012; Schneider et al., 2012), a freely available graphical user interface-based software, commonly used for image analysis in biological and medical contexts and students in section A could choose to follow the same handout as students in section B. The instructors explained to students how digital images can be treated as matrices of numbers, how to manipulate image brightness and contrast and how to use binary thresholding to isolate image features (**Figure 3B-D**). While our students used the MATLAB curve fitting toolbox to fit the data to the model, other relevant open-source packages in Python or R can be used as alternatives. Depending on availability of time and prior student knowledge, one might want to discuss curve fitting methods more broadly.

To increase the quality and size of students’ final data set without dramatically increasing the time needed for the module, the laboratory instructor prepared and mounted samples ahead of time for the subsequent sessions in weeks 3 and 4 (after all groups have prepared their own samples in the second week), so that students could focus on imaging and collecting data. The third and fourth weeks of the module were dedicated to the collection of more polished data, which was shared amongst the whole class to increase the collective sample size. Each student then analyzed the entire class’s data set independently.

### Presentation

Students were provided with resources on how to present experimental data, make figures and draft a report that contained their findings, comparisons to the literature and their interpretation of the data. While the rest of the lab module was a group effort, students were required to do this last part on their own. Once students had drafted their reports, they were asked to review the work of two of their peers. Peer review was conducted double-blind. We first discussed how to peer-review others’ work, emphasizing the importance of being both thorough and compassionate/constructive. 10% of the lab module’s participation points were based on peer-reviewing other students’ reports. In the Supplemental Material we provide a summary of how points were assigned to the different lab module components. The students then received their two reviews, so that they could incorporate the feedback and submit a final report. This exercise accomplishes several goals – it exposes students to the peer review process and helps them learn both, how to provide and how to receive feedback. It gives students a chance to think critically about material they are reading. It also exposes students to different ways of presenting and interpreting data.

## **IV.** Results and Discussion

### A. Experimental Results

Based on the final written reports, students largely confirmed the observations in Carter et al. that claimed the rate of mouth opening is consistent among *Hydra* despite variation in maximum mouth area between individuals. Students were able to successfully fit their recorded data to Eq. 1, the modified logistic equation, where *A(t)* is the normalized area of the mouth as a function of time, *t* is the time from the initiation of the opening process, *a* is the lower asymptote, *b* is the upper asymptote, *c* is the inflection point, and *d* is the rate of mouth opening (Carter et al., 2016), with R^2^ values > 0.90.

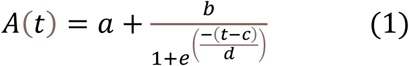

*a, b*, and *c* are experimentally constrained and correspond with initial mouth opening area, maximum mouth opening area, and 50% maximum mouth opening area. Once curves are normalized to the maximum opening, a is expected to be equal to 0 and b to be equal to 1. Parameter *d* is related to the length of time in which most of the mouth opening occurs and is inversely proportional to the rate of opening (**Figure 4**). Thus, faster openings correspond to smaller *d* values.

**Figure 4.**
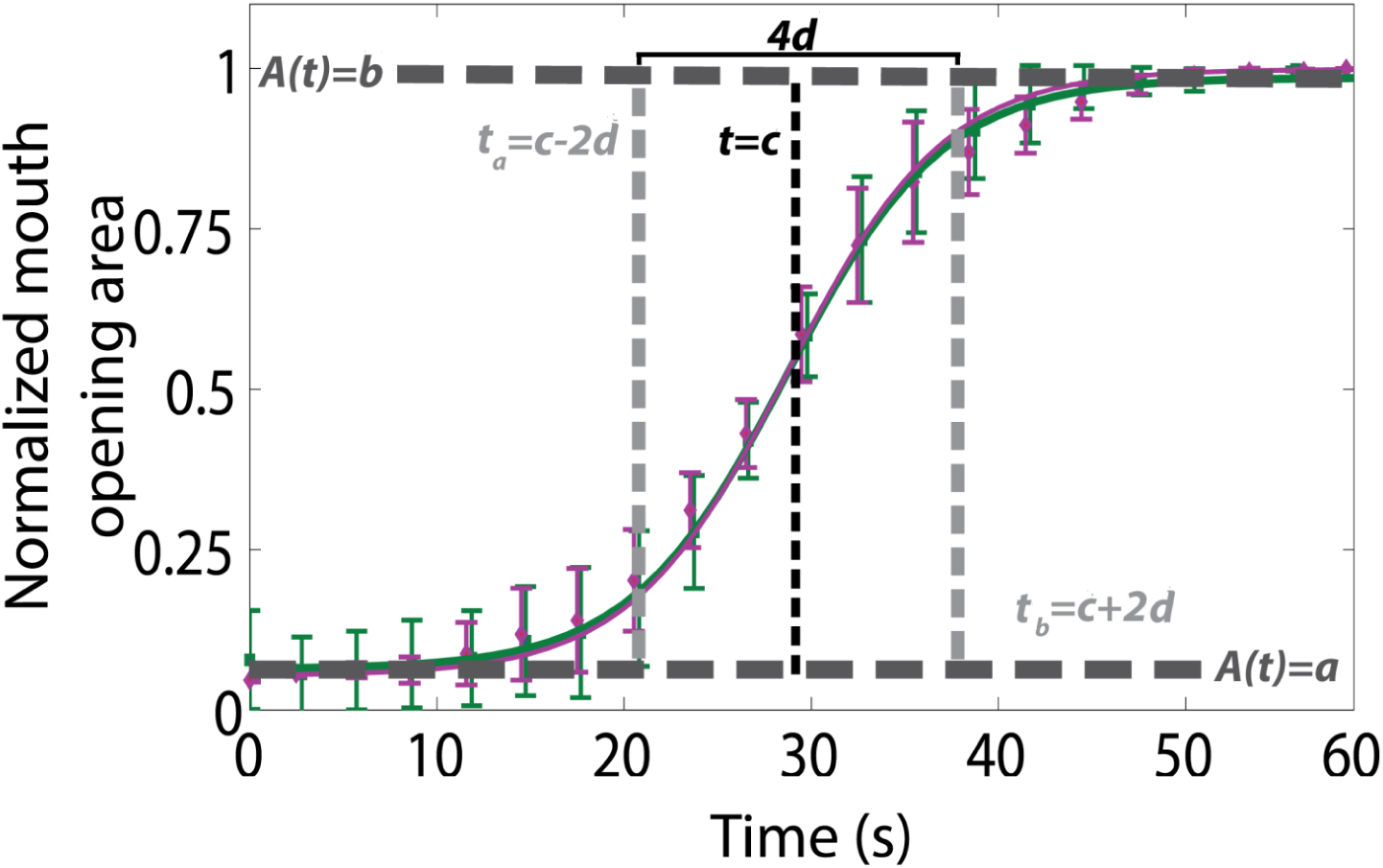
Definition of fit parameters. The normalized mouth opening area (A(t)) was fit according to Eq. 1. Data shows mean and standard deviations of mouth opening for the two epithelial layers, with ectoderm in green and endoderm in magenta. Figure reprinted from Carter et al. (Carter et al., 2016) with permission from authors.

Students found that their raw data collapsed to the characteristic sigmoidal shape after normalizing the opening area by the maximum area for each opening sequence (**Figure 5A, B**), as observed in the published paper. For some data, the mouth opening was more gradual, wherein the mouth first opened a small amount then paused before opening wider. This can be seen in **Figure 5A**, wherein one of the curves is longer and has a ‘bump’ before the typical S-shaped curve. For data that was suboptimal, students truncated the data as needed and set a to 0 and b to 1 instead of having them be fit parameters, to improve the quality of the fit.

**Figure 5.**
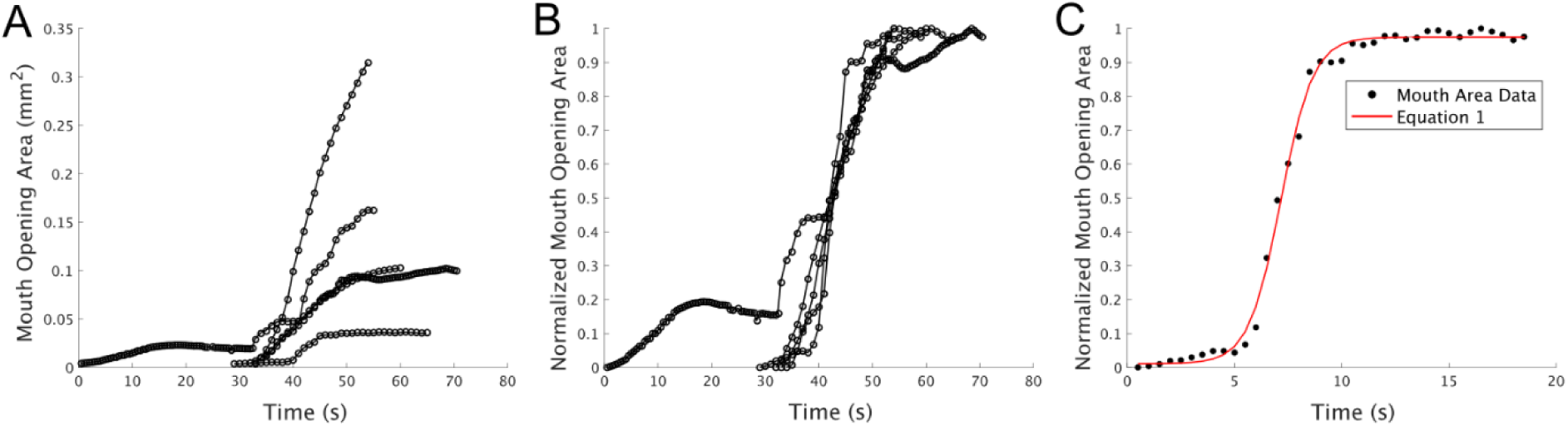
Example data of *Hydra* mouth opening dynamics from a student generated using MATLAB. (A) Raw (*N=5*) data aligned at the point in time at which the mouth opening reached 50% of its maximum value. (B) Normalized mouth opening data. (C) Student generated normalized *Hydra* mouth opening curve fitted to modified logistic equation (eq. 1). The data follows the expected S-shape for mouth opening. Because students independently decided the frame rate of their recordings, data was recorded at either 1fps or 2 fps. Since mouth area cannot be detected before opening, recordings do not start at 0 seconds.

An example of student reported *d* values based on the class data for individual mouth openings within a 95% confidence interval is (1.86, 4.94; n=5), which was much wider than the 95% confidence interval of (4.00,4.40; n=19) reported in Carter et al. However, the confidence intervals for the rate of mouth opening between student recordings and published data overlapped. Justifications for inconsistencies between the observed and published data included the following: differences in *Hydra* strains (Carter et al. used WM *Hydra*, whereas students used both WM and tricolor *Hydra* (Wang et al., 2022) due to animal availability) and differences in using spontaneous versus chemically induced mouth opening and type of inducer. Students used primarily quinine hydrochloride to induce the *Hydra* feeding response and only observed few spontaneous openings whereas Carter et al. analyzed primarily spontaneous openings and some that were induced by using reduced glutathione.

Common challenges students faced during their data analysis included using pooled image data from the class that had inconsistent or missing labels. Some students were unsure of their videos’ frame rate, which would affect the *d* value they calculated. This taught the students the importance of standardized naming conventions and detailed notes on experiments. Overall, students largely suggested that their observed chemically induced mouth opening occurred at a faster rate than the spontaneous mouth openings used in Carter et al.

### B. Student Confidence and Comfort

This module aimed to improve student aptitude for performing authentic interdisciplinary research by providing experience with live biological samples, fluorescent microscopy, image analysis, computation, and with reading, writing, and discussing scientific papers. The results of the student survey show that the successes of module were significant in these areas. **Figure 6A** shows the students’ self-reported comfort with various skills aligned with these goals and how this comfort changed after the completion of the module. Of the elements for which comfort was significantly improved (working with live animals, fluorescent microscopy, image analysis, programming, and scientific writing), four of the five categories showed the lowest incoming levels of comfort. We see the greatest increase in comfort (and the most statistically significant) around image analysis. We speculate that these gains may be attributed to the inclusion of authentic and interdisciplinary content, and to the inquiry-based approach to data analysis, which enabled students to more thoroughly engage with the computational techniques at hand. We also note that image analysis and programming show the lowest initial levels of comfort, reflecting the low incoming proficiency that students reported with programming (**Figure 2C**), suggesting that their lack of confidence is correlated with a lack of training and exposure. This result supports the claim that this module addresses shortcomings in the traditional laboratory curriculum, by building comfort in categories where the students were initially most apprehensive and inexperienced.

**Figure 6.**
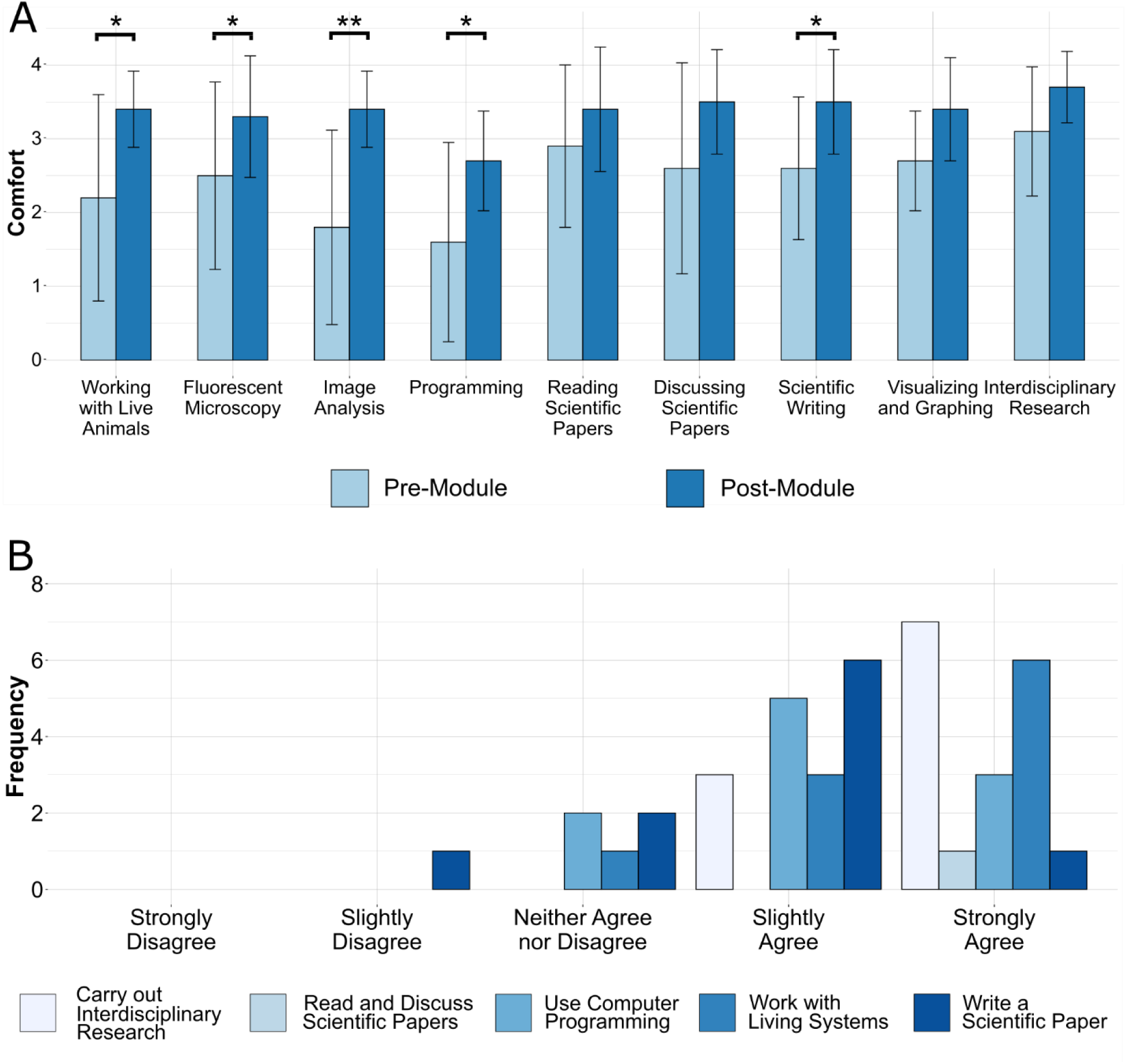
Self-reported comfort and confidence with lab module skills (n=10). (A) Student-reported comfort with various elements of the lab experience before (light bars) and after (dark bars) the *Hydra* mouth opening module. The score on the vertical axis represents an average comfort level, where 0 corresponds to “Very Uncomfortable” and 4 corresponds to “Very Comfortable”. Error bars represent the standard deviation. Asterisks indicate that the change in comfort from before to after the module for a given element is statistically significant, as calculated by a paired-sample sign test (one asterisk indicates p < 0.05, two asterisks indicate p < 0.01). (B) Stacked histograms of students’ agreement with the statement “The *Hydra* mouth-opening module made me more confident in my ability to [blank]”.

While students reported higher levels of comfort with reading scientific papers than with writing or discussing them (2.9 vs 2.6 and 2.6) before the module, their reported levels of comfort after the module were essentially equal (3.4 for reading, 3.5 for discussing and writing). This result, coupled with the significance of the increased comfort with scientific writing, suggests that the module meaningfully improves student comfort with writing scientific papers. Additional research would be required to identify how each of the individual writing-based reforms in this module impacted student comfort and could possibly be achieved by interviewing students as to what they think this could be contributed to.

Students were also asked about how their confidence with the learning goals of this module changed (**Figure 6B**). All student respondents to the survey (n=10) reported that this module made them more confident in their ability to read and discuss scientific papers and to carry out interdisciplinary research, and a majority reported confidence gains for the three other learning goals. While no direct assessment of student aptitude with each learning goal before and after the module was made, the marked increase in self-reported confidence shows the value of this module to empower students to engage in the research process. Moreover, 60% of students surveyed found the fact that the central paper was first-authored by an undergraduate to be inspiring, citing it as a source of engagement, interest and confidence. Specifically, in their open-ended response, one student said that “the *Hydra* mouth opening paper was inspiring because it served as an example of an undergraduate student (like myself and my classmates) having the opportunity and skills to make scientific contributions.” Another student said that “the fact that an undergraduate student wrote the paper made me more interested and engaged. It made me feel inspired and like I could be capable of doing this unit.”

#### Optional Advanced Exercises: Curve fitting, viscoelastic models of biological tissues

In our implementation of the lab module, students with a background in programming were encouraged to write their own code to analyze the image data. Building upon this first step, one could expand this lab module to teach classical image processing, including denoising images, thresholding binary images and segmentation. Students can then try extracting features of the mouth shape beyond its area, such as symmetry, circularity etc. Alternatively, students who do not have a background in programming can learn these same operations in ImageJ. They can further try to build an automated pipeline for image processing by using the ‘Record’ feature in ImageJ that generates ImageJ Macro code, based off Java language, for the steps students perform using the ImageJ graphical user interface. The students can then make minor tweaks to the generated code and directly run it in ImageJ to analyze mouth opening movies. For a more biophysical focus, students could be introduced to the concept of viscoelasticity in tissues and try to extract tissue relaxation times by fitting the latter half of the mouth opening area curve to an exponential function, as done in the Carter et al. paper. To further explore the mechanical behavior of viscoelastic tissues in response to forces, students could be asked to simulate a single spring dashpot system subject to an external force. Students can start by writing down the equation of motion for a spring dashpot system subject to an external force. They can then use numerical ordinary differential equation (ODE) solvers to observe how the spring dashpot system deforms in response to a constant external force or a sharp short-lived impulse/kick. In both cases, students should be encouraged to analyze the qualitative response of the system to these forces and test how changing the spring stiffness and/or dashpot viscosity affect the response time of the system. This is also a good opportunity to introduce the notion of viscoelastic relaxation time. Then, students can be asked to simulate other external force profiles – sinusoidal forces, exponentially decaying forces and so on and analyze the response of the system. Links should be made to how the spring dashpot system models soft tissue and the different force profiles model a range of mechanical conditions that different soft tissue experience. Students can be encouraged to think about what kinds of force profiles might generate a deformation like what they observe in mouth opening. Student can also try to further expand first to a linear chain of spring dashpots and then to a 2D network, see e.g. (Goel et al., 2024). Note that these exercises can become a semester long stand-alone module on modelling biological systems.

## V. Conclusion

This undergraduate *Hydra* mouth opening biomechanics laboratory module teaches fundamental skills, such as time-lapse microscopy, image analysis, programming, critical reading of scientific literature, and basics of scientific writing and peer-review. By using a research paper first-authored by an undergraduate student as the basis of this module, undergraduates can identify with the research, feel empowered, and thus are motivated to figure out how to do the research themselves. By offering two sections with different computational requirements, the module is broadly accessible to all students with introductory level biology and some classical mechanics knowledge. The additional exercises and the suggestions for learning opportunities that we discuss can be expanded to stretch this module to more weeks or to engage more advanced students.

## Use of human subjects

Approval was obtained from the Institutional Review Board (IRB-FY24-25-19) of Swarthmore College.

## Author Contributions

EMSC developed the lab module and designed and executed its implementation in the pilot described herein. JP was a student participant and produced the sample data shown in the results.

SH and EMSC developed the student survey and EMSC administered the survey. SH and EMSC analyzed survey results. TG and EMSC designed the advanced exercises. SH, TG, and EMSC wrote the initial draft of the manuscript. SH, TG, JP generated the figures. All authors edited the final draft.

## Acknowledgments

The authors have no conflicts of interest to report. This work was funded by NSF Grant 2102916 and Swarthmore College. The funders had no role in the design and conduct of the study; in the collection, analysis, and interpretation of the data; and in the preparation, review, or approval of the manuscript. The authors thank lab instructor Stacey Miller for her help with lab preparation, co-teaching the module, and discussions, Hannah Poon for providing the *Hydra* pinch response images, and Dr. Ben Geller for his comments on the manuscript.

