## Supplemental Information for "You can do it!–Using published undergraduate research on *Hydra* mouth opening to train undergraduates"

**Supplemental Methods**

**1. Week by week overview of the lab module and breakdown of activities:**
Below is a summary of the activities and goals of the module. Students were provided with a similar overview, informing them of the learning goals and objectives for each week.

<u>Week 1: Paper discussion</u>
At the beginning of the lab, the instructor gives a short 15-20min overview of this module and introduces the key biological and physical concepts that will be covered. Students arrive at the lab having already read the paper but are given another 30min to re-familiarize themselves and integrate the material the instructor covered in the introduction. They do this by filling in a worksheet, which is provided as a separate word document. After 30min, the instructor encourages students to discuss the worksheet with a partner and revise answers as needed/reach consensus. Then the whole class comes together to discuss the paper as a group. The worksheets are submitted for grading (thoughtful completion) to the instructor prior to the next lab session.

<u>Week 2: Acquaintance with experiment and equipment</u>
The instructor gives a short introduction to the equipment and *Hydra* mouth opening experiment. Students start with intact *Hydra* and go through the procedure of preparing *Hydra* head samples for imaging and attempt to take mouth opening data, following the provided step-by-step protocol below.

<u>Week 3: Take mouth opening data (heads are prepared ahead of time by lab instructor)</u>
Students apply the skills they learned in the previous week to collect mouth opening data and start the analysis, as outlined below.

<u>Week 4: Analyze data</u>
Students work on data analysis. Sample data are provided by the instructor if the students were not able to get data the previous week. Sample data also allow for remote instruction of the module, if needed.

<u>Week 5: Finish analysis and report writing</u>
Students continue to work on their analysis and start generating their figures and writing their reports. Scientific data presentation and good figure making is discussed are discussed as a group.

<u>Week 6: Peer review</u>
The instructor gives a short introduction to peer review and provides guiding documents for the review process to each student. We used a modified version of the materials developed by the Science Education Resource Center (SERC) at Carleton College

(https://serc.carleton.edu/sp/library/peerreview/tips.html) that included a few additional aspects, such as the confidentiality of the peer review and an emphasis on figure assessment. After discussing the document, students perform double-blind reviews of two of their peers' papers and submit their reviews at the end of the lab time. The instructor checks the reviews for completion and shares them with the students.

Week 7: Report submission
Students submit their final report and a rebuttal letter addressing the peer review comments for grading.

**2. Sample Preparation Protocol (as provided to the students)**
1. Pour your linalool solution (1mM) provided in the 15ml tube into the provided glass petri dish.

2. Use the glass pipet+bulb to transfer 5 animals into the glass petri dish. We will use animals that have not been fed in 2-3 days.

3. Keep the animals in linalool for 5-10 minutes to anesthetize them. The body column should extend.

4. Pinch the body column with tweezers. If the linalool is effective, the body column will swell only near where you pinched. This means your *Hydra* are ready to be cut.

5. Under the dissecting microscope, use the scalpel to cut the head at the base of the tentacle ring (as close to the tentacle ring as possible).

6. Transfer the head and the body column into a 100mm recovery dish (label!) filled with ~25ml of HM, transferring as little linalool as possible.

7. Let the animals recover from the anesthetic. It takes ~15-30 minutes for the effects of linalool to wear-off. You can confirm this by pinching the body column. An untreated *Hydra* contracts when pinched.

8. Put 1 strip of double-sided tape on either side of the glass slide, parallel to the narrow side, to create a small square well – that is slightly smaller than the 22x22mm glass cover slips you will use (**Figure S1**). Cut off excess tape with a razor blade on the cutting board.

9. Take the decapitated head and place it on the slide using the dissection scope in a drop of HM. Make sure the head is facing UP since you will be using an upright fluorescence microscope to image. Use forceps or a hair loop (**Figure S2**) as needed to ensure the head is oriented correctly and that no tentacles are in the way. If you got too much liquid on your slide, use a kimwipe to remove the excess HM without displacing the head.

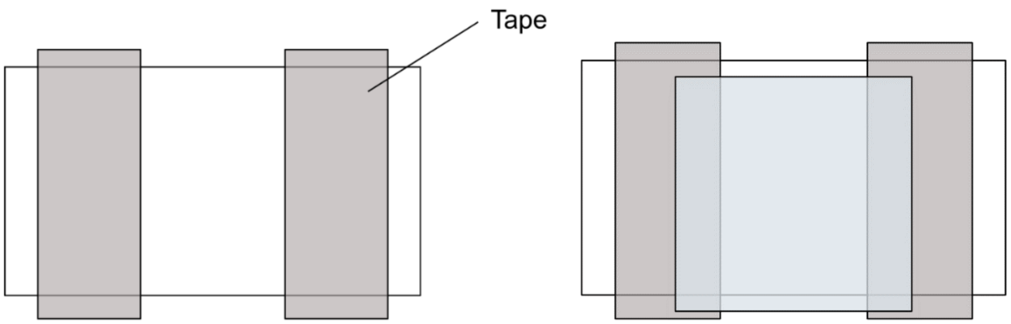


**Figure S1: Tunnel slide preparation.** Left: Tape (dark grey) on glass slide, spaced slightly smaller apart than the size of the glass cover slip that will be used. Right: Finished tunnel slide with cover slip (light blue) on top. Note the "lip" on the top. *Instead of placing the 22x22 coverslip with its side parallel to the larger coverslip, you can rotate it to a 45 degree angle and then place it on top of the tape. This would ensure that you always have access to the chamber.*

10. Gently place a coverslip (22mm x 22mm) on the slide as shown in **Figure S1** or at a 45 degree angle. Use the forceps to add the coverslip, you can fix one edge onto one side of the tape and let the opposite edge fall on the other piece of tape. It is important to ensure that the coverslip is flush with the glass slide on one side so that there is a "lip" on the other side of the glass slide where solutions can be flushed in. Now you have created a 'tunnel slide' that allows you to flush liquid in from one side. If there are air bubbles in your tunnel slide, carefully add extra HM with the P200 so that the entire slide is full, and the sample is hydrated.

11. The tunnel slide can now be imaged on the fluorescence microscope - make sure to tape the slide down before flushing in chemical solution so that the slide does not move around during flushing. Place the slide onto the microscope stage with the coverslip facing up.

#### Hair Loop Protocol

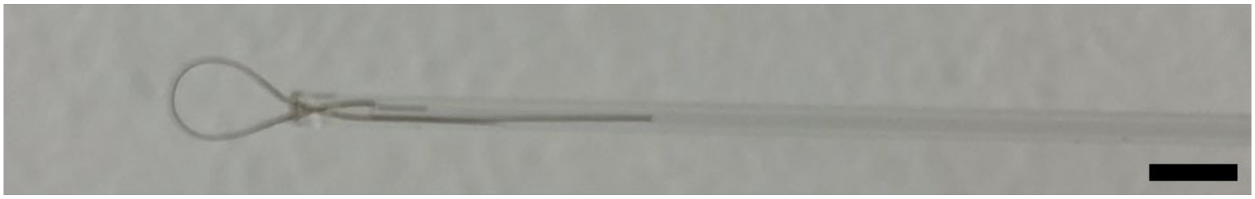

**Figure S2: Hair Loop.** Note the relative sizes of the hair strand, loop, and capillary tube. Scale bar: 3mm. Clear nail polish is used to seal the hair loop into the capillary tube.

1. Obtain a single small hair (preferably a darker colored hair – it is easier to see and stiffer)
2. Obtain a capillary tube (ideally close to a 25 µL capillary tube).
3. Push both ends of the hair into the capillary opening until it makes a small loop (**Figure S2**). If the hair is too long to do this easily, it can be cut shorter.
4. Seal with nail polish.
5. (If necessary) cut the capillary tube to a comfortable length.

### 3. Data Acquisition and Image Analysis

We used a USB 3 camera attached to a Nikon Ci-L compound light microscope to acquire images. A frame rate of 2 frames/sec provides sufficient temporal resolution for recording mouth opening but faster frame rates will yield better temporal resolution, especially during the fast opening period. Any software allowing for time series image acquisition can be used. After or before starting imaging, it is important to take an image of a micrometer with the same image settings being used to capture mouth opening data. This would provide the information necessary to convert the pixel dimensions in the image to a real-life length scale (**Figure S3**).

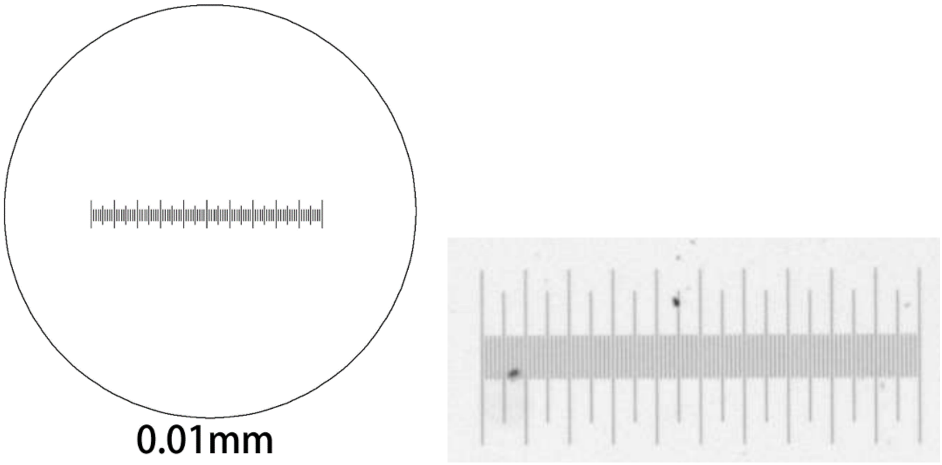


**Figure S3**. Micrometer images: Each tick is 10 microns. The distance between two large ticks is 100 microns.

**ImageJ analysis of mouth opening videos (as provided to the students)**

Step 1 – Reading in the data

Drag and drop the images folder that contains the data into the ImageJ toolbar. If you just want to view the image sequence and are not going to be making any permanent changes, click "**Open as Virtual Stack**".

Since we only want frames in which mouth opening occurs, the start frame should be the frame where the mouth is still fully closed, and the ending frame should be a frame where it is maximally open. Use **Image>Duplicate** to duplicate this part of the movie into a sperate window. Select "**Duplicate stack**" and enter the correct start and end frame.

Using the scale information obtained by imaging the micrometer, convert the image sequence from pixels to mm. To do this, select the line tool and draw a line over the known length. Convert pixel units to a standard unit of length by clicking **Analyze>Set Scale**. Enter the length corresponding to the line drawn on the image in "**Known distance**" box and change "**Unit of length**" from pixel to the chosen standard unit of length.

Step 2 – Isolating the mouth

The raw data images may be too dark to see the mouth opening. Change this by going to **Image>Adjust>Brightness/Contrast**. Make sure that the dark mouth has good contrast with the surround tissue. Click "**apply**" to make the change once satisfied with the image.

Crop the region of interest (ROI) tightly around the mouth (**Figure S4**). Use the rectangle tool and draw a box around the maximum mouth opening in the end frame and use **Image>Duplicate>Duplicate stack** to generate the cropped image stack.

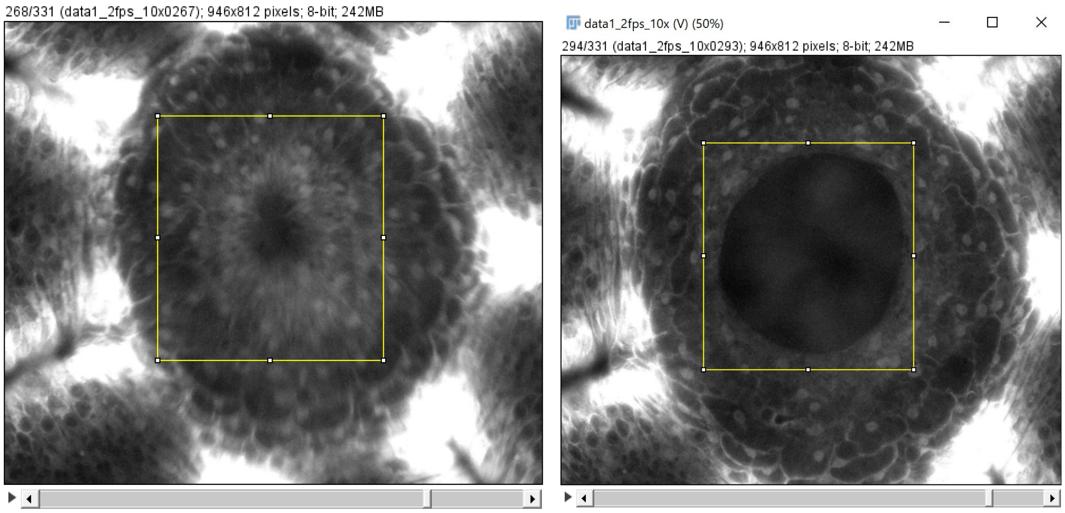


**Figure S4.** Image cropping. Use **Image > Duplicate** instead of cropping; this way you keep the original stack.

To isolate the ROI from the rest of the tissue, binarize image into mouth area and the background through thresholding. Use **Image>Adjust>Threshold**, unselect "**Dark background**" and "**Calculate threshold for each image**", and click "**Apply**". The correct threshold needs to be found empirically through adjusting the sliding bars (**Figure S5**).

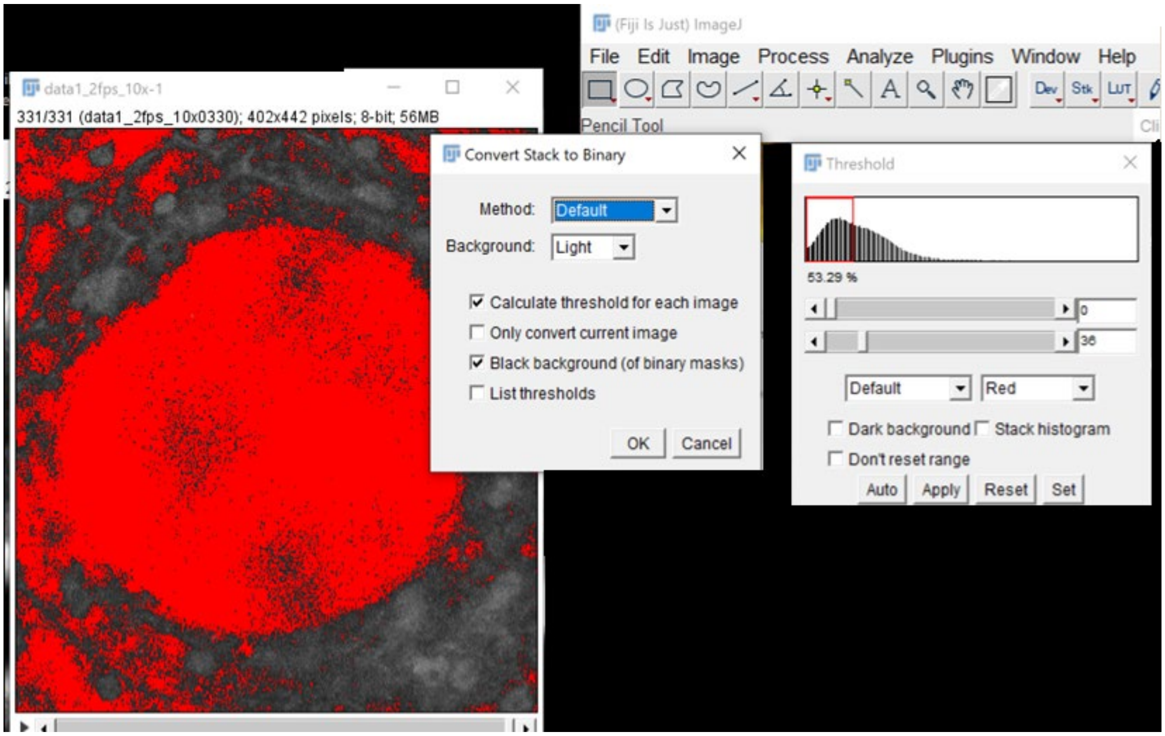


**Figure S5.** Example of the thresholding process

Step 3 – Measuring the mouth area in every frame

To specify measurements for area analysis, click **Analyze>Set>Measurements**, and check **Area** and **Stack position**. This will tell you whether you have more than 1 object per frame. To extract mouth area from every frame, select **Analyze>Analyze Particles**. Select "**Display results**", "**Clear results**", and "**Show outlines**".

*You likely need to set a minimum particle size to eliminate any noise you might have in your image. If you don't know where to start, run the analysis without any filter and see how large the mouth area is - you can find it by looking at the labeled objects in the 'Display outlines' window and looking up the object number in the 'Results' window. Then you can re-run the analysis with a filter that is right below that minimum mouth area size. It might take a few trials to get this setting right.*

After the analysis is finished, the 'Results' window will automatically open. The 'Slice' indicates the number of particles detected in each frame. If there are more than one particle counted or zero particles counted at any frame, re-run the function with different minimum and maximum particle size or manually delete frames. Save the results as .csv or .txt file to import these values into MATLAB and continue analysis.

Step 4 – Curve Fitting

Create a MATLAB script in the same folder you stored your results. Use the **readtable** function to read in your .csv file as a table. Store both the frame and area values as an array through the **table2array** function. Convert frames to time by multiplying the frames by your frame rate. Normalize the mouth area by the maximum mouth opening area.

Open the MATLAB Curve Fitting Toolbox by typing **cftool** in the command window. Create a new fit for each mouth opening and choose matching time and normalized area arrays in "Select Data". In "Custom Equation", type in Eq. 1, as shown in **Figure S6** and check the values of the fit parameters.

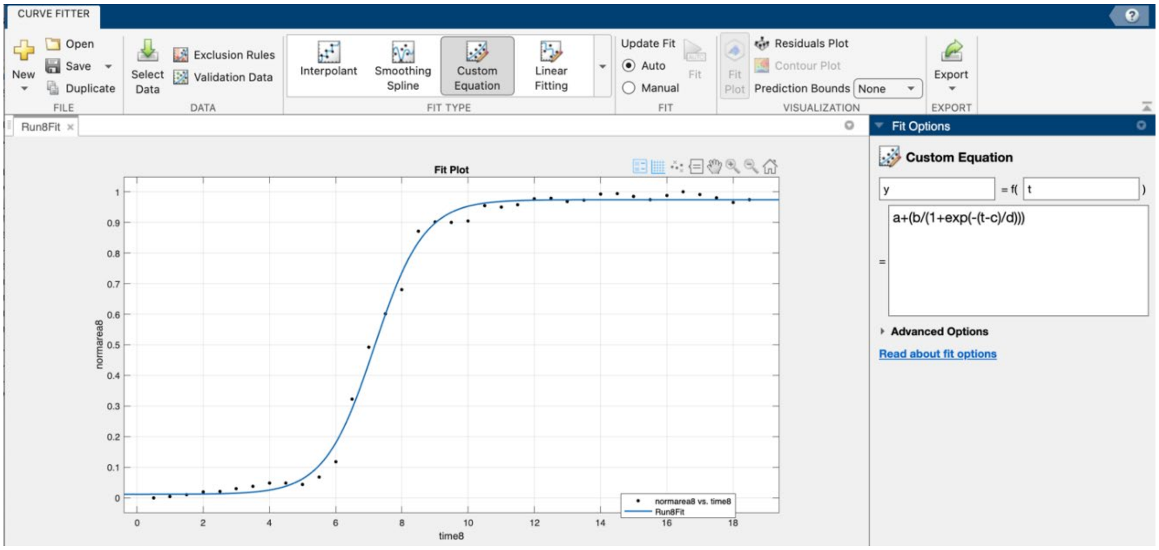


**Figure S6.** Example of curve fitting mouth opening data with MATLAB Curve Fitting Toolbox.

**4. Short Guidelines for Writing the Report (as provided to the students)**

The elements below are expected to be part of your final report for this lab module.
Different (good) ways of doing these things and showing your data are possible.
Remember: This work must be your own!

The report should be maximum 5 pages long and contain:

- A figure (schematic, still images, a mix) explaining the problem at hand: Hydra mouth / mouth opening. This figure should accompany a short text (max 2-3 paragraphs) explaining the project in your own words (like a mini-introduction)
- A figure summarizing your analyzed data
- A figure showing your fit of the data with the phenomenological S-shaped curve
- A narrative / summary of the results
- A discussion of the results and comparison to the literature
- References as appropriate

**5. Point distribution and grading**

The grade breakdown for this module was as follows:
Participation: 25%
Experimental/analysis: 25%
Write-up (final report with rebuttal letter): 50%
The participation points included participation in discussions, quality of worksheet reflections, effectiveness of group work, and the two peer reviews (10%). The experimental/analysis part included both the quality of data generated and the quality of protocol execution, note taking, and overall diligence, taking into consideration that behavioral work with live biological specimen can be variable. The write-up portion of the grade included the final report and rebuttal letter.

**Supplemental Data**

Four example raw imaging data sets are provided as separate .zip files, to allow for a remote version of this laboratory, if necessary. In addition, we have included a compressed .csv file containing 5 sheets, each corresponding to a separate mouth opening, which can be shared with students for analysis practice. Each sheet contains the time (s), area ($mm^2$), and the fps of one mouth opening event.

**Supplemental Movie**

**Movie S1:** Student sample data of *Hydra* mouth opening. Single-channel movie showing a transgenic Watermelon animal undergoing quinine hydrochloride induced mouth opening. using a transgenic watermelon animal. The move is recorded at 2 fps and played at 10 fps. Scale bar: 100 μm.
